# Clickable methionine as a universal probe for labelling intracellular bacteria

**DOI:** 10.1101/784371

**Authors:** Sharanjeet Atwal, Suparat Giengkam, Yanin Jaiyen, Heather Feaga, Jonathan Dworkin, Jeanne Salje

**Affiliations:** Public Health Research Institute, Rutgers University of New Jersey, USA; Mahidol Oxford Tropical Research Unit, Bangkok, Thailand; Department of Microbiology & Immunology, College of Physicians and Surgeons, Columbia University, USA; University of Oxford, UK

## Abstract

Despite their clinical and biological importance, the cell biology of obligate intracellular bacteria is less well understood than that of many free-living model organisms. One reason for this is that they are mostly genetically intractable. As a consequence, it is not possible to engineer strains expressing fluorescent proteins and therefore fluorescence light microscopy – a key tool in host-pathogen cell biology studies – is difficult. Strain diversity limits the universality of antibody-based immunofluorescence approaches. Here, we have developed a universal labelling protocol for intracellular bacteria based on a clickable methionine analog. Whilst we have applied this to obligate intracellular bacteria, we expect it to be useful for labelling free living bacteria as well as other intracellular pathogens.

## 1. Introduction

Obligate intracellular bacteria cause a range of human and veterinary diseases around the world. The two main orders of obligate intracellular bacteria are the Rickettsiales and Chlamydiales. Chlamydiales cause sexual- and aerosol-transmitted diseases in humans and are the leading cause of non-congenital blindness worldwide. The Rickettsiales are spread by arthropod vectors and most have animal reservoirs. Rickettsial species cause a wide range of human diseases including typhus (*R. prowazekii)*, Rocky Mountain Spotted Fever (*R. rickettsii*) scrub typhus (*Orientia tsutsugamushi*), Anaplasmosis (*Anaplasma*) and Ehrlichiosis (*Ehrlichia*) [1-4], whilst *A. marginale* causes disease in cattle [5]. The Rickettsiales *Wolbachia* is not known to cause disease but is a widely distributed endosymbiont of arthropods and nematodes [6].

Fluorescence light microscopy is an important tool for understanding host-pathogen cell biology, especially in the case of obligate intracellular bacteria where a visualization of the interactions between bacteria and host is indispensable for an understanding of their interactions. Most obligate intracellular bacteria remain genetically intractable [7, 8] and therefore fluorescent protein-based approaches to labeling bacteria are not possible. Immunofluorescence based approaches have been extremely powerful and are currently the main tool for labelling obligate intracellular bacteria. However, the Rickettsiales are a very diverse order and antibodies generally need to be generated specifically for each organism. Even where genetic tools are available, this needs to be repeated for any new environmental and clinical isolates limiting workflow and throughput. For this reason, we have been developing universal tools to label obligate intracellular bacteria.

We recently reported the use of a panel of fluorescent reporters that could be used to label bacteria for live cell imaging [9]. In the current work we have built on this work by developing protocols for a methionine-based probe. In addition to being used to delineate intracellular bacteria, this probe reports on the metabolic activity of bacteria under study.

Here, we have used a clickable, non-toxic methionine analog probe (L-Homopropargylglycine, HPG) which readily incorporates into newly synthesized proteins to label a range of obligate intracellular bacteria from the order Rickettsiales [10]. The methionine derivative is conjugated to an alkyne (or azide) moiety and is added to growing bacterial cells. Cells are fixed, and then the incorporated methionines are conjugated to an azide (or alkyne) coupled fluorophore using a copper catalyzed click reaction (Fig. 1). This allows metabolically active bacteria to be visualized by fluorescent microscopy.

**Fig 1.**
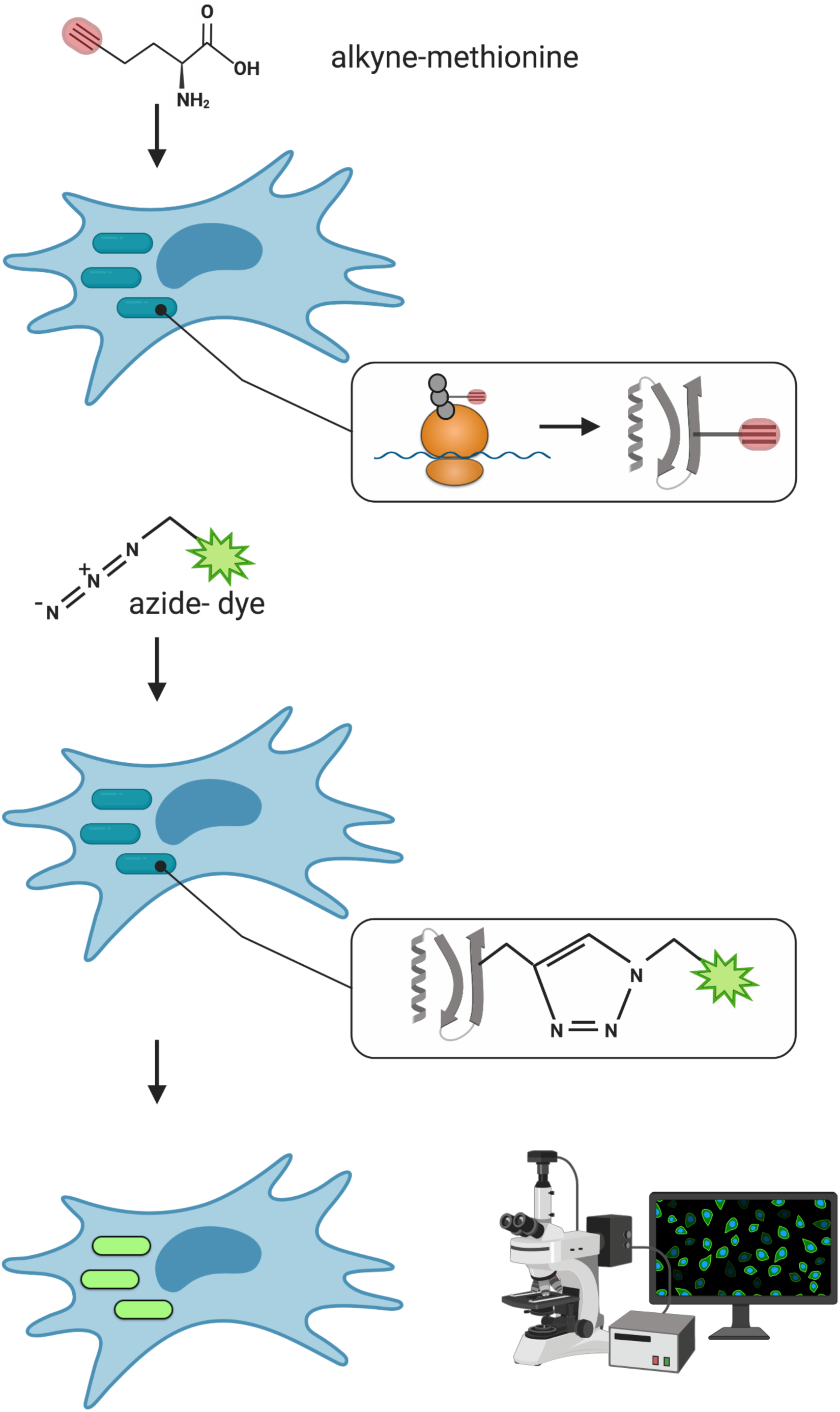
Schematic overview of HPG labelling of intracellular bacteria. Created with BioRender.

## 2. Materials and methods

### Growth of bacteria and cell lines

The following bacterial strains were used: *Orientia tsutsugamushi* strain Karp, *Rickettsia canadensis* (gift from Nancy Connell, Rutgers University), *Anaplasma marginale* strain Oklahoma 291, *Wolbachia pipientis* endosymbiont of *Aedes albopictus, Ehrlichia chaffeensis* strain AR (all gifts from Ulrike Munderloh, University of Minnesota) and *Anaplasma phagocytophilum* strain HGE1 (gift from Thomas Bakken, University of Minnesota).

Macrophage-like DH82 cells (ATCC CRL-10389) were grown in 25cm^2^ flasks with Eagle’s Minimum Essential Medium (EMEM) (Sigma, M0325, USA) with 10% heat inactivated FBS at 37°C and 5% CO_2_. Human leukemia HL-60 cells (ATCC CCL-240) were grown in 25cm^2^ flasks with Iscove’s Modified Dulbecco’s Medium (IMDM) (ATCC 30-2005) with 10% heat inactivated FBS at 37°C and 5% CO_2_. *Aedes albopictus* clone C6/36 cells (ATCC CRL-1660) were grown in 25 cm^2^ flasks with Eagle’s Minimum Essential Medium (EMEM) (Sigma, M0325, USA) with 10% heat inactivated FBS at 28°C and 5% CO_2_. L929 cells (ATCC CCL-1) were grown in RPMI 1640 Medium with HEPES (Thermo Fisher Scientific, 22-400-071, USA) supplemented with 10% heat inactivated FBS (Thermo Fisher Scientific, 16140071, USA) in 25cm^2^ flasks at 35°C and 5% CO_2._ Kidney epithelial Vero cells (ATCC CCL-81) were grown in RPMI 1640 Medium with HEPES, supplemented with 10% heat inactivated FBS in 25cm^2^ flasks at 37°C and 5% CO_2_.

The bacterial strains were grown in the following cell lines: *Orientia* in L929 cells (as shown previously [11]), *R. canadensis* and *A. marginale* in Vero cells, *A. phagocytophilum* in HL-60 cells, *E. chaffeensis* in DH82 cells and *Wolbachia* in C636 cells.

Chloramphenicol was used at 150 µg/ml and was added to infected host cells at the same time and for the same duration as the methionine probe. Cycloheximide was used at 40 µg/ml and added to infected host cells for 4 hours prior to addition of the methionine probe.

### Methionine labelling and click chemistry

Probes Click-IT™ L-Homopropargylglycine (HPG) (Thermo Fisher Scientific, C10186, USA) or L-Azidohomoalanine (Click Chemistry Tools, 1066-25, USA) were added to growing bacteria in host cells at 50µM in methionine-free DMEM culture media (Thermo Fisher Scientific, 21013024, USA) or methionine supplemented media (Thermo Fisher Scientific, 11965092, USA. After 30 minutes, the probe and culture media were discarded, and the cells were fixed with 4% PFA for 10 minutes at room temperature. Cells were washed with 3X PBS and permeabilized with 0.5% triton X100 for 10 minutes on ice. Click chemistry was performed using the Click-iT™ Cell Reaction Buffer Kit (Invitrogen, C10269, USA) with an azide or alkyne Alexa Fluor 488/594 (AF 594 Alkyne, 1297-1 or AF 488 Picolyl Azide 1276-5, Click Chemistry Tools, USA) for 1 hour in the dark. Cells were washed with PBS 3x and covered with mounting media (20mM Tris, pH 8.0, 0.5% N-propyl-gallate and 90% glycerol. All imaging was performed using either Zeiss LSM700 (Carl Zeiss, Jena, Germany) or Leica DMi8 laser scanning confocal microscopes (Leica Microsystems, Wetzlar, Germany).

## 3. Results and discussion

### HPG specifically labels bacterial protein synthesis

To confirm that the methionine probes were specifically labelling bacterial protein synthesis, chloramphenicol was added to infected host cells. Chloramphenicol targets the bacterial ribosome and inhibits protein synthesis. Bacterial labelling was inhibited by chloramphenicol treatment indicating specific incorporation of HPG into nascent proteins (Fig. 2).

**Fig 2.**
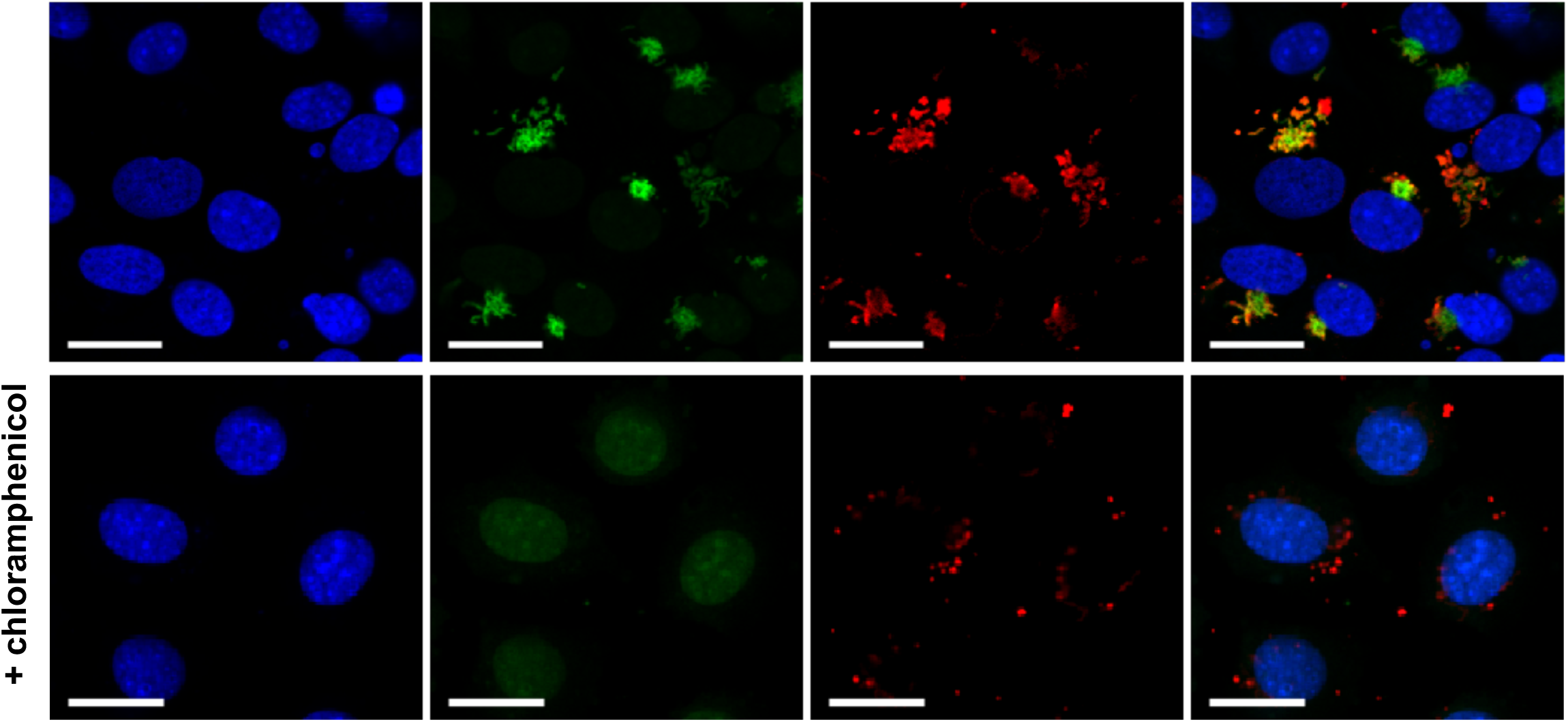
Bacterial ribosome inhibitor chloramphenicol blocks nascent protein synthesis labelled by clickable methionine-alkyne (HPG) probe. *Orientia tsutsugamushi* were grown in L929 host cells in the presence of HPG for 30 mins, cells were fixed, and HPG was reacted with an azide-conjugated fluorophore to reveal cells actively undergoing protein synthesis (green). Bacteria are counterstained in red (TSA56 antibody) and host cell nuclei are shown in blue (DAPI). Scale bar = 20μm

### The signal from HPG-labelled bacteria can be detected over the signal from host cell protein synthesis

Both bacteria and host cells undergo protein synthesis and therefore both will incorporate HPG that is present in the growth media. We found that when incubating with the probe for 1-3 hours the signal from intracellular bacteria could be readily detected against the background of host cells (Fig. 3). However, host cell background could be further reduced using a specific inhibitor of eukaryotic protein synthesis, cycloheximide. A concentration of 40ug/ml reduced the host signal but did not affect host morphology. This may be beneficial in those cases where the bacterial signal is weak or when a very strong signal to noise ratio is required.

**Fig 3.**
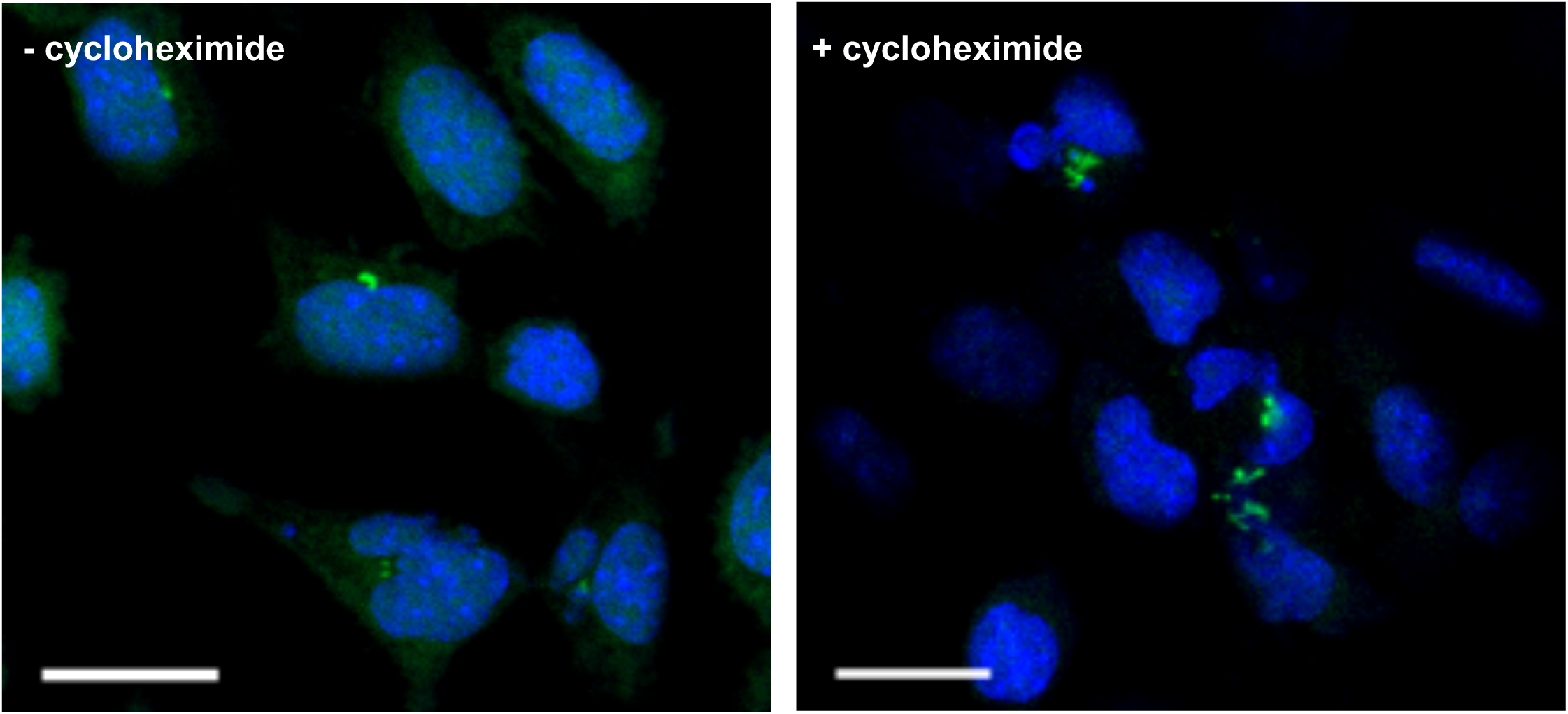
Eukaryotic protein inhibitor cycloheximide can reduce the background from host cells. *Orientia tsutsugamushi*-infected L929 host cells were treated with 50ug/ml cycloheximide for 4 hours. Methionine HPG probe was added to infected cells for 30mins, cells were fixed, and HPG was reacted with an azide-conjugated fluorophore to reveal cells actively undergoing protein synthesis (green). Host cell nuclei are shown in blue (DAPI). Scale bar = 20μm

### Use of methionine-free media can improve signal-to-noise

Mammalian cell culture media usually contains a supplement of methionine (e.g. RPMI contains 0.015g/L and DMEM contains 0.03g/L). This will compete with the clickable methionine analog for incorporation into growing polypeptides. We tested the effect of this by comparing HPG labelling in media with and without unlabelled methionine. Whilst bacterial labelling could be clearly resolved in both cases there was an increase in signal when incubation was performed using methionine-free growth media (Fig. 4).

**Fig 4.**
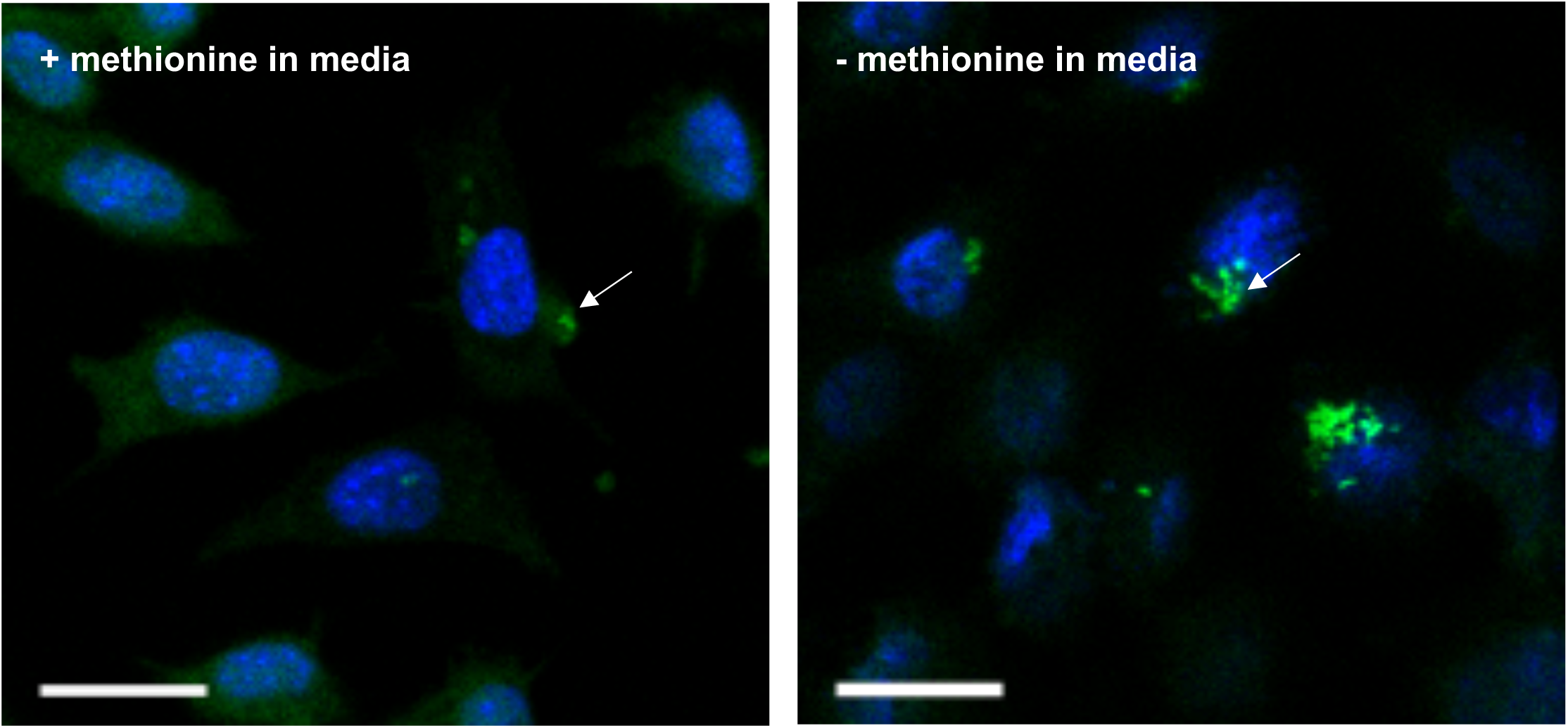
Methionine-free media can improve intracellular bacterial labelling. *Orientia tsutsugamushi* cells were grown in L929 host cells with HPG for 30mins in media supplemented with methionine (left) or methionine-free media (right). Cells were fixed, and HPG was reacted with an azide-conjugated fluorophore to reveal cells actively undergoing protein synthesis (green). Host cell nuclei are shown in blue (DAPI). White arrows show HPG-labelled bacteria. Scale bar = 20μm

### Using HPG labelling to image a panel of Rickettsiales obligate intracellular bacteria

We tested the ability of HPG to label a diverse group of 6 obligate intracellular Rickettsiales bacterial species (*Anaplasma marginale, Ehrlichia chaffeensis, Anaplasma phagocytophilum, Wolbachia, Orientia tsutsugamushi, Rickettsia canadensis*) (Fig. 5). Despite differences in host cell type, replication cycle and generation time all could be labelled successfully using the same protocol. Obligate intracellular bacteria are dependent on host cells for provision of nutrients, therefore any bacteria outside host cells were metabolically inactive and not labelled with the methionine probe.

**Fig 5.**
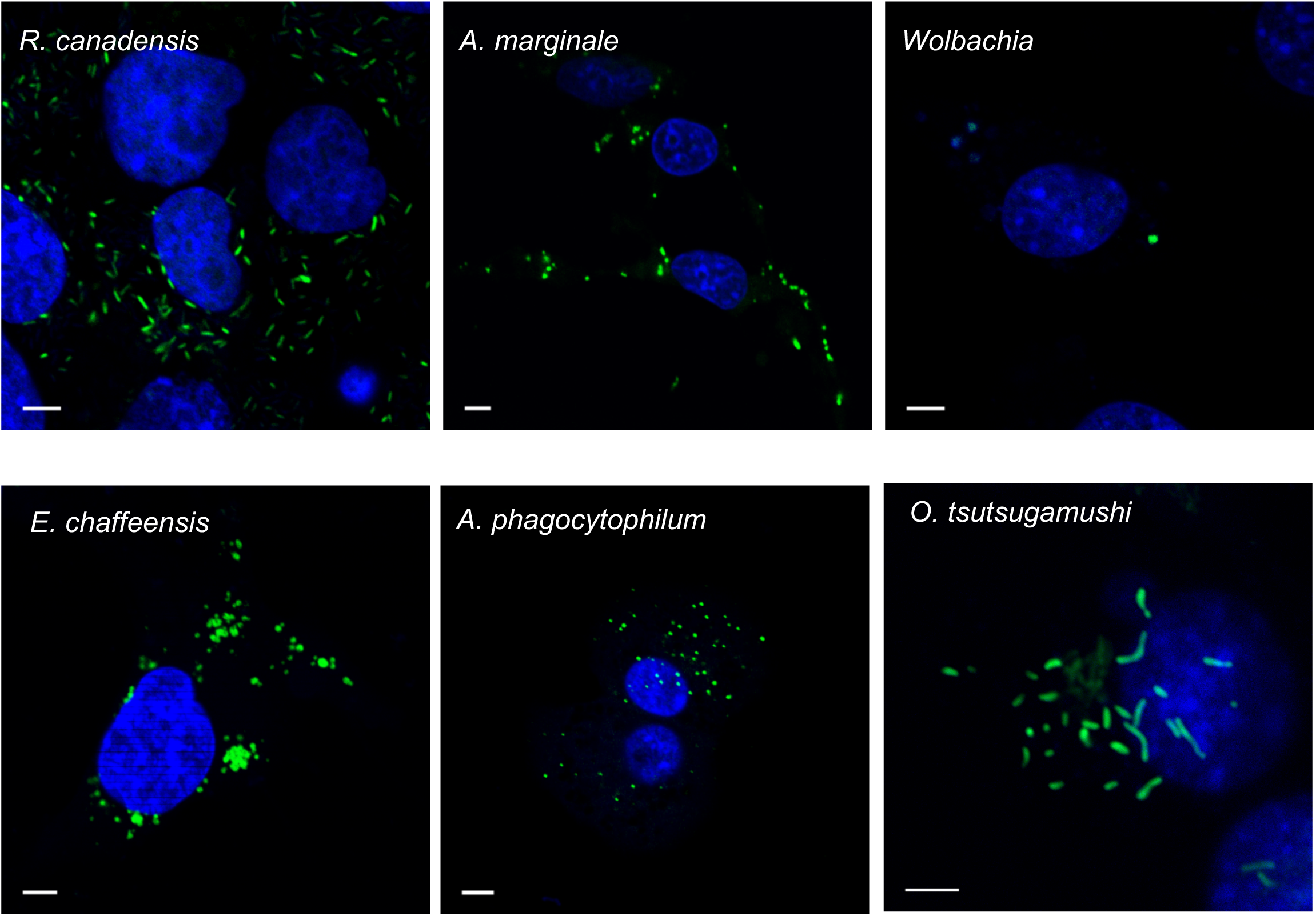
HPG can label Intracellular Rickettsiales species. The methionine probe HPG was added to infected cells for 30 minutes prior to fixation. HPG was reacted with an azide-conjugated fluorophore to reveal cells actively undergoing protein synthesis (green). Host cell nuclei are shown in blue (DAPI). Scale bar = 5μm

## 4. Conclusion

Taken together, these results show that a clickable methionine probe can be used to fluorescently label obligate intracellular bacteria. This probe can be used as a universal visual label for pathogens for which other tools are not available. In addition to delineating bacterial organisms it can be used to probe for metabolic activity within bacterial populations.

## 5. Acknowledgements

HF was supported by NIH F32GM122266; JD was supported by R01 GM114213, R21AI135427, and is a Burroughs-Welcome Investigator in the Pathogenesis of Infection Disease. JS was supported by a Royal Society Dorothy Hodgkin Research Fellowship DH140154

## References

1. Luce-Fedrow, A., et al., A Review of Scrub Typhus (Orientia tsutsugamushi and Related Organisms): Then, Now, and Tomorrow. Trop Med Infect Dis, 2018. 3(1).

2. Fang, R., L.S. Blanton, and D.H. Walker, Rickettsiae as Emerging Infectious Agents. Clin Lab Med, 2017. 37(2): p. 383–400.

3. Battilani, M., et al., Genetic diversity and molecular epidemiology of Anaplasma. Infect Genet Evol, 2017. 49: p. 195–211.

4. Saito, T.B. and D.H. Walker, Ehrlichioses: An Important One Health Opportunity. Vet Sci, 2016. 3(3).

5. Kocan, K.M., et al., Antigens and alternatives for control of Anaplasma marginale infection in cattle. Clin Microbiol Rev, 2003. 16(4): p. 698–712.

6. Miller, W.J., Bugs in transition: the dynamic world of Wolbachia in insects. PLoS Genet, 2013. 9(12): p. e1004069.

7. McClure, E.E., et al., Engineering of obligate intracellular bacteria: progress, challenges and paradigms. Nat Rev Microbiol, 2017. 15(9): p. 544–558.

8. Salje, J., Orientia tsutsugamushi: A neglected but fascinating obligate intracellular bacterial pathogen. PLoS Pathog, 2017. 13(12): p. e1006657.

9. Atwal, S., et al., Live imaging of the genetically intractable obligate intracellular bacteria Orientia tsutsugamushi using a panel of fluorescent dyes. J Microbiol Methods, 2016. 130: p. 169–176.

10. Beatty, K.E., et al., Selective dye-labeling of newly synthesized proteins in bacterial cells. J Am Chem Soc, 2005. 127(41): p. 14150–1.

11. Giengkam, S., et al., Improved Quantification, Propagation, Purification and Storage of the Obligate Intracellular Human Pathogen Orientia tsutsugamushi. PLoS Negl Trop Dis, 2015. 9(8): p. e0004009.

